# Female-biased vascular smooth muscle cell gene regulatory networks predict MYH9 as a key regulator of fibrous plaque phenotype

**DOI:** 10.1101/2025.03.28.645955

**Authors:** R Noah Perry, Graham Lenert, Ernest Diez Benavente, Angela Ma, Nicolas Barbera, Michal Mokry, Dominique P V de Kleijn, Menno P J de Winther, Manuel Mayr, Johan L M Björkegren, Hester M den Ruijter, Mete Civelek

## Abstract

Atherosclerosis, a chronic inflammatory condition driving coronary artery disease (CAD), manifests in two primary plaque types: unstable atheromatous plaques and stable fibrous plaques. While significant research has focused on atheromatous plaques, recent studies emphasize the growing importance of fibrous plaques, particularly in females under 50 years of age, where erosion on fibrous plaques significantly contributes to coronary thrombosis. The molecular mechanisms underlying sex differences in atherosclerotic plaque characteristics, including vascular smooth muscle cell (VSMC) contributions, remain understudied. Therefore, we utilized sex-specific gene regulatory networks (GRNs) derived from VSMC gene expression data from 119 male and 32 female heart transplant donors to identify molecular drivers of fibrous plaques. GRN analysis revealed two female-biased networks in VSMC, GRN_floralwhite_ and GRN_yellowgreen_, enriched for inflammatory signaling and actin remodeling pathways, respectively. Single-cell RNA sequencing of carotid plaques from female and male patients confirmed the sex specificity of these networks in VSMCs. Further sub cellular phenotyping of the single-cell RNA sequencing revealed a sex-specific gene expression signature within GRN_yellowgreen_ for VSMCs enriched for contractile and vasculature development pathways. Bayesian network modeling of the GRN_yellowgreen_ identified *MYH9* as a key driver gene. Indeed, elevated MYH9 protein expression in atherosclerotic plaques was associated with higher smooth muscle cell content and lower lipid content in female plaques, suggesting its involvement in fibrous plaque formation. Further proteomic analysis confirmed MYH9’s upregulation in female fibrous plaques only and its correlation with stable plaque features. These findings provide novel insights into sex-specific molecular mechanisms regulating fibrous plaque formation.

## Introduction

Atherosclerosis, the underlying cause of coronary artery disease (CAD), is a chronic inflammatory disease characterized by the accumulation of lipids, inflammatory cells, and extracellular matrix (ECM) components within the arterial wall^1^. Substantial progress has been made in understanding the pathophysiology of atherosclerosis; however, significant gaps remain. Pathology studies have described two dominating plaque morphologies with unique pathological mechanisms. Atheromatous plaques, unstable and prone to plaque rupture, are characterized by a thin fibrous cap and a large necrotic core consisting of accumulated lipids and inflammatory cells. Fibrous plaques, stable and prone to plaque erosion, are characterized by an intact, thick fibrous cap rich in smooth muscle cells (SMCs) and ECM^2,3^. In the past decades, most attention has been focused on understanding plaque rupture hallmarked by large atheroma and plaque hemorrhage^4^. However, it is estimated that 40% of coronary thrombosis in sudden cardiac death cases involving CAD is associated with plaque erosion. Further, in females under the age of 50, this percentage increases to ∼75-80%^2,5^. These studies highlight the importance of further understanding the risk of both stable and unstable plaques and the contributing factors of sex differences in the underlying pathogenesis of atherosclerotic disease.

Systems biology approaches identifying gene regulatory networks (GRNs) are an effective tool in studying the complexity of common disorders such as CAD^6,7^. These approaches postulate that networks of genes, rather than individual expression changes or linear pathways, define complex physiological and pathological processes. Highly connected key driver genes are essential for the activity and impact of GRNs on CAD^8^. Recently, such methods have been employed to investigate GRNs of atherosclerotic tissue between males and females^9,10^.

Identifying key drivers of female-specific GRNs has increased mechanistic understanding of the regulation of female-biased atherosclerotic lesions. However, such studies generate GRNs using expression profiles from a heterogeneous population of cells, and there may be significant differences between individual cell types^11^. The female-biased atherosclerotic networks pointed to SMC plasticity as an important phenomenon. Indeed, SMC content and phenotypic state play a critical role in defining the characteristics of fibrous versus atheromatous plaques. SMCs migrate from the medial layer into the intima to form the fibrous cap, but their phenotypic plasticity enables a range of behaviors with different impacts on disease^12^. In addition to known cellular composition differences between male and female plaques^3^, sex hormone studies highlight critical roles for both estrogen and testosterone in SMC phenotypic transitions^13–16^.

Despite the well-established sex differences within SMCs in the clinical presentation of CAD and associated risk factors, our understanding of the underlying genetic and molecular mechanisms that drive these differences remain largely unexplored. Understanding SMC-specific GRNs may, therefore, better pinpoint individual key driver genes contributing to sex differences in males and females associated with plaque erosion.

We generated sex-specific GRNs from gene expression data of vascular smooth muscle cells (VSMCs) isolated from 119 male and 32 female heart transplant donors from distinct genetic ancestries^17^. We used network-based preservation statistics to identify intra-network topology differences across male and female GRNs^18^. To investigate VSMC contributions to plaque erosion, we focused on the subset of female-specific networks with unique VSMC gene-to-gene interactions. We prioritized GRNs with previous evidence of female-biased connectivity as well as genetic and biologic signatures associated with VSMC pathology in atherosclerosis. This approach led to the selection of two female-specific GRNs with sex-biased genetic regulation enriched for genes involved in inflammatory signaling (GRN_floralwhite_) and vascular and actin remodeling (GRN_yellowgreen_) – each relevant to known characteristics that differ between plaque erosion and rupture phenotypes^19,20^. Utilizing single-cell RNA sequencing data from the Athero-Express Biobank of patients undergoing carotid endarterectomy^21^, we validated the female-biased nature of both networks but only found additional evidence supporting the rewiring of actin cytoskeleton remodeling processes between males and females in the context of disease (GRN_yellowgreen_). We then employed a Bayesian network approach and key driver analysis and identified *MYH9* as the key driver gene of GRN_yellowgreen_. Finally, we incorporated both *in vitro* and *in vivo* phenotypic data specific to atherosclerosis phenotypes. This provided evidence that MYH9 regulates a female-specific, fibrous-associated VSMC signaling process and is directly associated with the high prevalence of fibrous plaques in females. This study provides new insights into the sex-specific molecular mechanisms driving plaque phenotypic differences.

## Methods

### Smooth Muscle Cell Culture and Gene Expression

Smooth muscle cell (SMC) gene expression dataset and donor characteristics have been described in detail elsewhere^17^. Briefly, we cultured aortic SMCs isolated from 6, 12, 64, and 69 individuals with East Asian, African, Admixed American, and European ancestries in complete media (containing 5% FBS) until 90% confluence. We then switched to either serum-free media for 24 hours to mimic the quiescent state of SMCs or continued to culture in complete media to mimic the proliferative state of SMCs^22^. Total RNA was extracted using the RNeasy Micro Kit (Qiagen) and the RNase-free DNase Set. RNA integrity scores for all samples, as measured by the Agilent TapeStation, were greater than 9, indicating high-quality RNA preparations.

Sequencing libraries were prepared with the Illumina TruSeq Stranded mRNA Library Prep Kit and were sequenced to ∼100 million read depth with 150 bp paired-end reads at the Psomogen sequencing facility. We trimmed the reads with low average Phred scores (<20) using Trim Galore^23^ and mapped the reads to the hg38 version of the human reference genome using the STAR Aligner^24^. We quantified gene expression by calculating the transcripts per million (TPM) for each gene using RNA-SeQC based on GENCODE v32 transcript annotations. In addition to protein-coding RNAs, we also measured the non-coding RNA since they have been shown to play significant roles in SMC biology^25^. We considered a gene as expressed if it had more than 6 read counts and 0.1 TPM in at least 20% of the samples. RNAseq data is available from GEO with the accession number GSE193817.

### Quantification of Atherosclerosis-relevant Cellular Phenotypes

Atherosclerosis-relevant cellular phenotypes capturing calcification, proliferation, and migration were previously quantified and characterized for the isolated aortic SMCs^17^. Briefly, to measure calcification, VSMCs were cultured in complete media until 90% confluence, then switched to either commercially available STEMXVivo human osteogenic media (R&D Systems) or complete media supplemented with inorganic phosphate to final concentration of 3.7 nM^26^ for 27 days before determining calcium content using the CPC kit according to the manufacturer’s instructions (ThermoFisher Scientific). Migration was continuously monitored for 15 minutes for 24 hours using electronically integrated 16-well Boyden chamber plates (ACEA Biosciences) with 8-um pores in the xCelligence Real-Time Cell Analysis Instrument (ACEA Biosciences) placed in a 5% CO2 humidified incubator maintained at 37C. Cells were seeded in the upper chamber containing serum-free media and monitored as cells migrated to the lower chamber containing either serum-free media (control) or serum-free media supplemented with 100 ng/mL PDGF-BB. For each donor, the migration response difference was measured by estimating the area-under-the-curve, determining differences in slopes, and calculating minimum and maximum cell index values between PDGF-BB and control media. Finally, to test proliferation, cells were cultured in serum-free media containing either 20 ng/mL TGF-B1, 10 ng/mL, or 25 ng/mL IL-1B for an additional 24 hours. Serum-free media without cytokines was used as control. BrDU labeling reagent was added during the last eight hours of cytokine treatment, and BrDU incorporation was determined using the BrDU cell proliferation assay kit (Cell Signaling). Relative proliferation was calculated with respect to a reference donor and proliferation in response to cytokine stimulus was calculated by the change in proliferation in each cytokine condition with respect to the serum-free media condition.

### Weighted Gene Co-expression Network Analysis

A gene module is a cluster of densely interconnected genes in terms of co-expression. Independent modules were generated for each of the four experimental conditions using female gene expression data from quiescent VSMCs, female gene expression data from proliferative VSMCs, male gene expression data from quiescent VSMCs, and male gene expression data from proliferative VSMCs separately as model inputs. The same input list of 11,300 genes was used for each sex-by-condition test. We used Iterative Weighted Gene Co-expression Network Analysis (iterativeWGCNA)^27,28^, which uses hierarchical clustering and an adjacency matrix, to identify gene modules. IterativeWGCNA follows the same principles as WGCNA but re-runs WGCNA iteratively to prune poorly fitting genes resulting in more refined modules than WGCNA. The adjacency matrix is defined as the similarity between the i-th gene and j-th gene based on the absolute value of the Pearson correlation coefficient between the profiles of genes i and j. Soft-thresholding powers are applied to the adjacency matrix to reduce the noise of correlations and create a network that resembles a scale-free graph representative of biological systems. Scale-free graphs are characterized by a power-law distribution where few hub nodes exist and new nodes prefer to connect with existing nodes^29,30^. To best control for potential differences in network topology due to set input parameters, the soft thresholding power was determined by choosing the lowest shared power across the four experimental conditions that yielded a log-log R^2^ scale-free model fit ≥ 0.8 (**Supplemental Figure 1**). Genes that are not assigned to any of the modules are designated to the grey module. Because these genes are not co-expressed, we did not consider them in our analyses. For each gene in a given co-expression module, we calculated the sum of the intramodular connectivity across all co-expressed genes. Genes within the top 10% of intramodular connectivity strengths were defined as hub genes.

### Network Preservation Analysis

We performed preservation analysis on constructed modules across both sexes and conditions. Female modules generated from the proliferative condition were tested for preservation against modules generated from the male proliferative condition. Similarly, female modules generated from the quiescent condition were tested for preservation against the male quiescent condition. To determine whether a network of genes is perturbed between sexes of the same VSMC culture condition, we studied modules whose connectivity patterns are not preserved between sexes as demonstrated by their module preservation statistics. For this analysis, we used the summary statistic, medianRank, implemented in the WGCNA R package as a composite module preservation statistic^31^. medianRank is a rank-based measure that relies on observed preservation statistics. medianRank is calculated as the mean of medianRank.density and medianRank.connectivity. Density is the mean adjacency (connection strength) across all nodes (genes) in the network. Connectivity is the sum of connection strengths with the other network nodes. To calculate medianRank.density and medianRank.connectivity, for each statistic 𝑎 in the reference network, we ranked modules in the test network based on the observed values 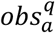 Thus, each module is assigned a rank 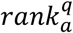 or each observed statistic. The median density and connectivity ranks are then calculated for each module, 𝑞, in the test network. A module with a lower medianRank exhibits stronger observed preservation statistics than a module with a higher median rank. We identified the least preserved modules by defining the modules scoring in the bottom 20th percentile of preservation (modules with the highest medianRank score).

### Pathway enrichment of co-expression modules

To interpret the biological significance of the 10 replicated female-biased co-expression modules, we performed enrichment analysis using Gene Ontology^32^ Biological Process and Molecular Function gene sets with the anRichment R package. We used the model input list of 11,300 genes as the background gene set and employed an FDR cutoff of 0.2 to signify enrichment.

### Coronary Artery Disease-associated gene list

Coronary artery disease-associated loci, identified in a Genome-Wide Association Study (GWAS), may contain several potentially causal genes for the disease. To capture all genes associated with a given loci, we employed results from five studies to curate a list of CAD-associated genes, which resulted in a prioritized list of 684 predicted causal genes^33–37^.

### Sex-specific genetic associations with atherosclerosis-relevant cellular phenotypes

Twelve atherosclerosis-relevant phenotypes related to calcification, proliferation, and migration were previously described for the cohort of VSMCs isolated from 151 multi-ethnic heart transplant donors^17^. For each of the genes in the two prioritized genetic regulatory networks (GRNs; floralwhite and yellowgreen), we first performed a series of two-way ANOVA tests to examine whether there was an interaction effect between sex and gene expression on cellular phenotype. Each of the 12 phenotypes was then ranked for each GRN according to the number of genes with a significant sex-by-gene expression interaction effect (nominal p ≤ 0.05). For the top-ranked phenotype associated with each GRN, Pearson correlations for each gene in the GRN and the phenotype were calculated using gene expression values from only females, only males, and all samples.

### Study Populations

The Athero-Express Biobank is an ongoing, prospective biobank study, collecting atherosclerotic plaques from patients undergoing carotid endarterectomy in two Dutch tertiary referral hospitals: University Medical Centre Utrecht and St. Antonius Hospital, Nieuwegein. The study design and inclusion criteria have been published previously^21^. We used previously published GRNs generated from gene expression data from advanced plaques of 158 age-matched females and males for comparisons with sex-specific data generated in this study^10^. Plaque samples of 46 patient plaques (20 females and 26 males) were used for scRNA-sequencing analysis. Untargeted proteomics was conducted on 200 plaques (51 female and 149 males) to study sex differences in atherosclerotic plaques at the protein level^38^.

The Stockholm-Tartu Atherosclerosis Reverse Network Engineering Task (STARNET) is a multi-tissue transcriptomic study composed of 600 patients who underwent a coronary artery bypass grafting procedure, of which 30% are female. Details of the study and its population have been described before^39^. We used previously published GRNs generated from gene expression data from atherosclerotic aortic root tissue of 160 age-matched females and males for comparisons with sex-specific data generated in this study^9^.

### Plaque Histopathology

As described previously^3,10,21^, the atherosclerotic plaque was processed directly after surgery. According to a standardized protocol, the plaque was divided into segments of 5-mm thickness along the longitudinal axis. The segment with the greatest plaque burden was subjected to histological examination. Semiquantitative estimation of the plaque morphology was performed at ×40 magnification for SMC content (α-actin) and amount of collagen (picrosirius red).

Histological plaque characteristics were scored as (1) no or minor staining or (2) moderate or heavy staining. The criteria for classification were defined as follows: for smooth muscle cells: (1) no or minor α-actin staining over the entire circumference with absent staining at parts of the circumference of the arterial wall; (2) positive cells along the circumference of the luminal border, with locally at least few scattering cells; and for collagen staining: (1) no or minor staining along part of the luminal border of the plaque; (2) moderate or heavy staining along the entire luminal border. In addition, smooth muscle cell content was scored as the percentage of the total plaque area with the specific staining by using computerized analyses to validate the semiquantitative analyses. The size of the lipid core was estimated visually as a percentage of the total plaque area with the use of H&E and picrosirius red stains, with a division into categories of <40% and >40% of the total plaque area based on the correlation of the lipid core size and plaque stability^40^. Plaques with<10%, 10–40%, and >40% fat were categorized as fibrous, fibro-atheromatous, and atheromatous, respectively. All plaque characteristics were scored and quantified with good intra- and interobserver reproducibility by two independent observers. Presenting symptoms and duplex stenosis were retrieved from patient charts.

### Single-cell RNA sequencing of atherosclerotic plaques

Single-cell RNA sequencing (scRNAseq) was performed on 46 plaques (20 females and 26 males) and yielded a total of 4,948 cells (1,962 female cells and 2,986 male cells). The process of preparing, sorting, and sequencing cells from plaques was described in detail previously^41,42^. scRNAseq data was processed in R-3.6.2 using the Seurat-3.2.2 R-package^43^. Mitochondrial genes and doublets were filtered out from the data, setting thresholds for unique and total reads per cell. Batch effects were corrected using SCTransform. Clustering was performed with 20 principal components, validated through a JackStraw analysis to ensure significant feature distribution. A clustering resolution of 0.8 was chosen to reflect meaningful biological groupings without overfitting. Sensitivity analysis showed minimal impact of clustering parameters on cell-type identification. Optimal clustering parameters were finalized after multiple iterations. Cell populations were annotated using differential gene expression analysis, and automated cell type classification was performed with SingleR (version 1.2.4), which compared cluster profiles to the BLUEPRINT reference dataset. SMCs were classified by expression of *ACTA2* and *TAGLN*. Subpopulations of SMCs were identified by further clustering using 10 principal components at a higher resolution, and distinct identities were assigned based on differential gene expression analyzed via enrichR-3.0^44^. The module score for differentially expressed genes was calculated using the addModuleScore function in Seurat^43^, providing an expression proxy for gene sets. Genes that were not represented in the scRNAseq were removed from this calculation.

### Bayesian Network Generation and Validation

Bayesian Networks (BN) were constructed on continuous expression data to infer directed, acyclic relationships between nodes in a module using BNLearn due to its improved performance for smaller module sizes^45,46^. Within BNLearn, we used the Hill-climbing algorithm (HC). HC employs a greedy search to generate acyclic graphs via random insertions, removals, and reversals of edges. HC was utilized to create a network via BNlearn’s boot.strength function, which employs nonparametric bootstrapping to infer edge weights and directions for every possible edge in the module. We used a bootstrap replicate value of 10,000. The edge weight represents the proportion of appearances of the edge in replicates compared to the total number of replicates, with a value of 1 denoting complete consistency across replicates.

Once strengths for all available edges were generated, a procedure to remove low-weight edges was performed. To ensure representative biological modeling within our trimmed network, we opted to set an edge removal threshold that would yield a scale-free topological distribution in the final graph. A scale-free distribution follows a power-law distribution, with few nodes having high edge connectivity and many nodes having lower edge connectivity^47^. To quantitatively define adherence to a power-law distribution in a network generated by a given elimination threshold, the R^2 value of a log(Frequency) vs log(Degree) graph was calculated. An R^2 value greater than .5 indicated the threshold weight yielded a graph structure more representative of a power-law distribution and, therefore, a scale-free topology. Further, a hypothesis test via BNlearn’s bootstrap_p function was performed to determine if a given edge removal threshold yielded a network that was not representative of a power-law distribution.

1000 simulations per threshold were performed. For each edge weight elimination threshold, the subsequent R^2 value was calculated, and the hypothesis test was performed. The lowest threshold greater than 0.4 that passed both criteria was chosen as the final network threshold. The final network was generated with this threshold value, having edges below the elimination threshold weight pruned from the original graph. All remaining edges were then considered unweighted within the final graph.

### Key Driver Analysis

To identify key regulators for a given regulatory network, we performed key driver analysis (KDA)^48^, which takes as input a set of genes (G) and a directed gene network (N). KDA first generates a sub-network N_G, defined as the set of nodes in N that are no more than h-layers away from the nodes in G. We first computed the size of the h-layer neighborhood (HLN) for each node in the reconstructed BN. For the given network N, *μ* was defined as the average size of the HLN. A score was added for a specific node if the HLN was greater than *μ* + 𝜎(*μ*). Total key driver scores for each node were then defined as the summation of all scores at each h-layer scaled according to h. The top 10% of key driver scores > 0 were defined as key driver genes.

### Untargeted LC-MS proteomics in atherosclerotic plaques

Untargeted proteomics was conducted on 200 plaques (51 female and 149 males) to study sex differences in atherosclerotic plaques at the protein level. Details regarding the protein extraction and quantification have been described in detail elsewhere^38^. In brief, samples underwent a 2-step protein extraction process, initially with a NaCl buffer to isolate loosely bound proteins, followed by a guanidine hydrochloride (GuHCl) buffer to solubilize mature ECM proteins, with both fractions stored at -80°C. GuHCL extracts were quantified using a Pierce BCA protein assay, and 20µg of protein from each sample was precipitated with ethanol for deglycosylation. A two-step deglycosylation process was employed forthe GuHCL extracts to remove glycosaminoglycans and other sugar monomers, preparing the samples for in-solution digestion with trypsin. The digested samples were then purified using a C18 cartridge system and analyzed by LC-MS using an untargeted proteomics approach on a nano-flow LC system. Spectra were collected with an Orbitrap mass analyzer, and data-dependent MS2 scan was performed for the top 15 ions. Data analysis involved a database search against the human UniProtKB/Swiss-Prot database with specific modifications and enzyme settings. Protein identification yielded 2,148 proteins, with ECM and related proteins categorized using Matrisome DB and further in-house selection. Protein abundances were filtered, normalized, and scaled, with missing values imputed using the KNN-Impute method, resulting in a final dataset of 1,499 proteins for further analysis^49^. Differential abundance analysis was performed using Limma^50^. We selected proteins that were differentially abundant between female and male fibrous plaques (p < 0.05, logFC > 0.3, AveExpr > 4). Differentially abundant proteins were enriched using clusterProfiler^51^.

## Results

### Generation of sex-specific gene expression modules in human VSMCs

We constructed sex-specific gene coexpression networks from RNA-sequencing data of aortic smooth muscle cells (SMCs) isolated from 119 male and 32 female heart transplant donors from distinct genetic ancestries. SMCs were cultured in quiescent or proliferative conditions to mimic different vascular conditions represented in atherosclerosis^17^. **Figure 1** summarizes the overall analysis workflow used in this study. For each sex and culture condition, we input 11,300 genes for weighted gene coexpression network analysis^27,28^ (WGCNA) to identify groups of genes with high topological overlap. We employed a soft-threshold power of 5 for each network condition to control for variations that could be introduced into the model (**Supplemental Figure 1**). This was the single shared value at which all networks were at or near a scale-free threshold without risk of over-fitting^30,47^. The female quiescent and proliferative SMC conditions resulted in 10,885 and 10,829 coexpressed genes segmented into 55 and 68 modules, respectively. The male quiescent and proliferative conditions resulted in 10,633 and 9,942 coexpressed genes segmented into 43 and 53 modules, respectively (**Supplemental Table 1**). The female datasets resulted in more modules, potentially due to smaller sample sizes.

**Figure 1:**
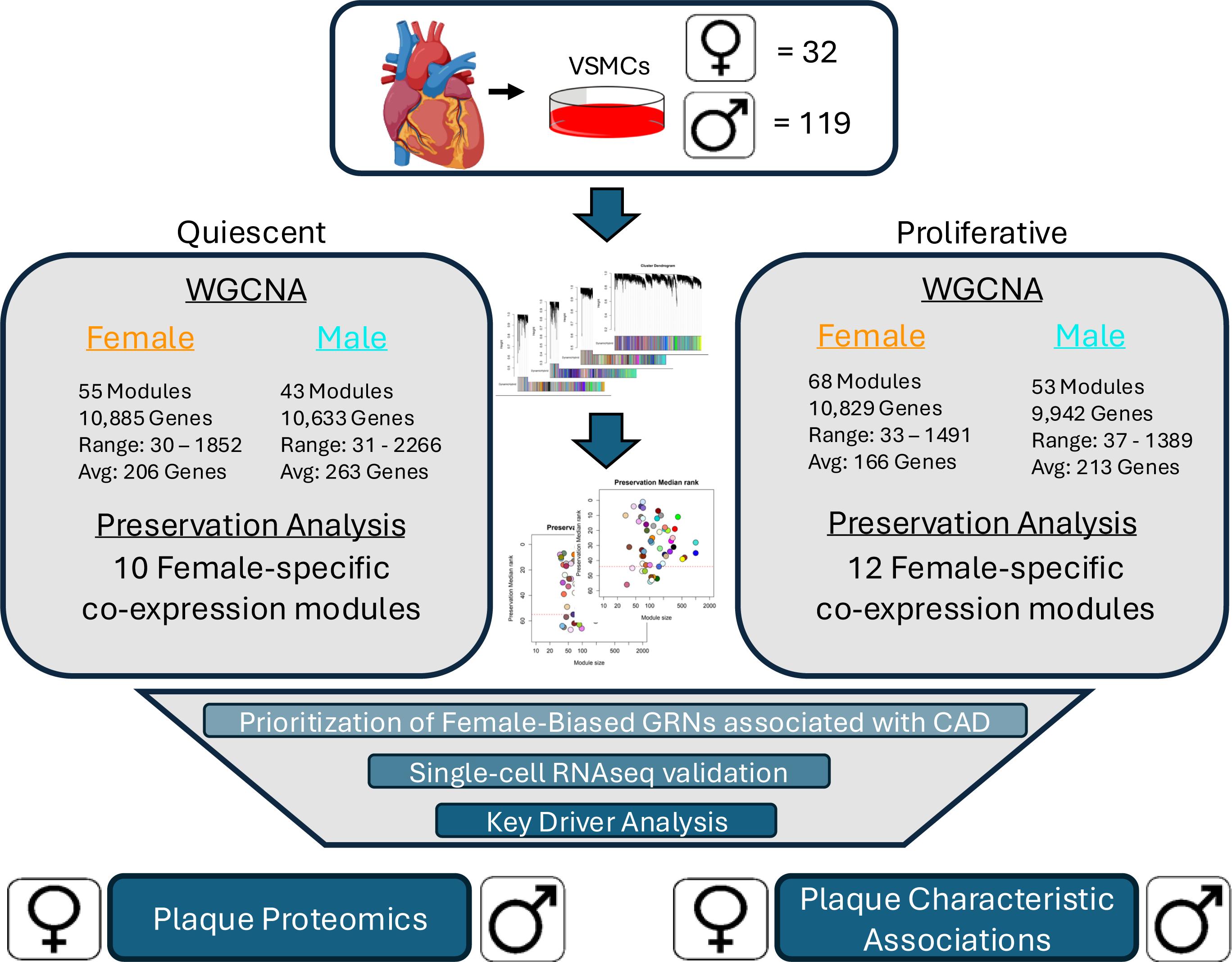
Schematic representation of the overall study design. Vascular smooth muscle cells (VSMCs) from the ascending aortas of 32 female and 119 male donors were cultured with and without FBS to mimic the quiescent and proliferative phenotypes of VSMCs. Gene expression was measured with RNA sequencing and used as input into WGCNA to generate sex-specific coexpression modules for each phenotypic condition. Network preservation analysis was performed between sexes for each phenotypic condition to rank modules based on preservation. Sex-specific modules were prioritized according to associations with CAD and sex-biased reproducibility across multiple datasets. Prioritized coexpression modules were then reconstructed into Bayesian networks to perform key driver analysis to identify key regulating genes of female-specific genetic architecture linked to CAD development. Proteomics data and plaque characteristics from patients undergoing carotid endarterectomy were used to link key driver genes to disease risk and plaque development.

To test the potential effects of using different sample sizes for males and females, we performed a sensitivity analysis on the quiescent and proliferative male datasets. We randomly subsampled 32 male samples five times and re-constructed coexpression networks. We observed an increase in the number of modules compared to the coexpression networks constructed from all 119 male samples, suggesting that sample size affects module membership (**Supplemental Table 2**). Therefore, we studied if these module differences resulted in limitations in capturing biologically meaningful information. For this, we performed pathway enrichment analyses across the modules using Gene Ontology Terminology (GO Terms)^32,52^, Kyoto Encyclopedia of Genes and Genomes (KEGG)^53,54^, and Hallmark^55^ pathways. Across all three pathway databases, modules generated from a smaller sample size (n=32) led to fewer enriched pathways detected (FDR ≤ 0.05). However, of these enriched pathways, 70% were reproduced within the pathway enrichment results from modules generated with 119 male samples (**Supplemental Table 2**). While some information may be missed when using 32 randomly selected male samples, this analysis suggests that there is enough power to detect groups of coexpressed genes enriched for biologically relevant information. Also, performing downstream analysis comparing the female and male networks may provide valuable insight in SMC pathophysiology.

### Prioritization of female-specific modules in human VSMCs

To prioritize female-specific GRNs to elucidate key regulators of plaque, we first identified female GRNs that were not preserved in the male GRNs. We assessed whether the 55 GRNs identified in quiescent female VSMCs were preserved in the 43 GRNs identified in quiescent male VSMCs. We performed a similar analysis for the proliferative female (68) and male (53) GRNs. Using a composite statistic of preservation from the WGCNA R package, medianRank^31^, we ranked the preservation of each GRN across sexes in each respective phenotypic state. A low medianRank score represents a highly preserved GRN expected to capture similar biological regulation and processes shared across sexes, whereas a high medianRank score represents a less preserved GRN expected to identify genes and biological pathways most likely to be specific to female VSMCs. Therefore, we identified the GRNs scoring in the bottom 20th percentile of preservation, leading to 10 female-biased GRNs in the quiescent condition and 12 female-biased GRNs in the proliferative condition **(Figures 2A,B)**. These GRNs contain 968 and 1627 genes in the quiescent and proliferative phenotypes, respectively **(Supplemental Table 3**).

**Figure 2:**
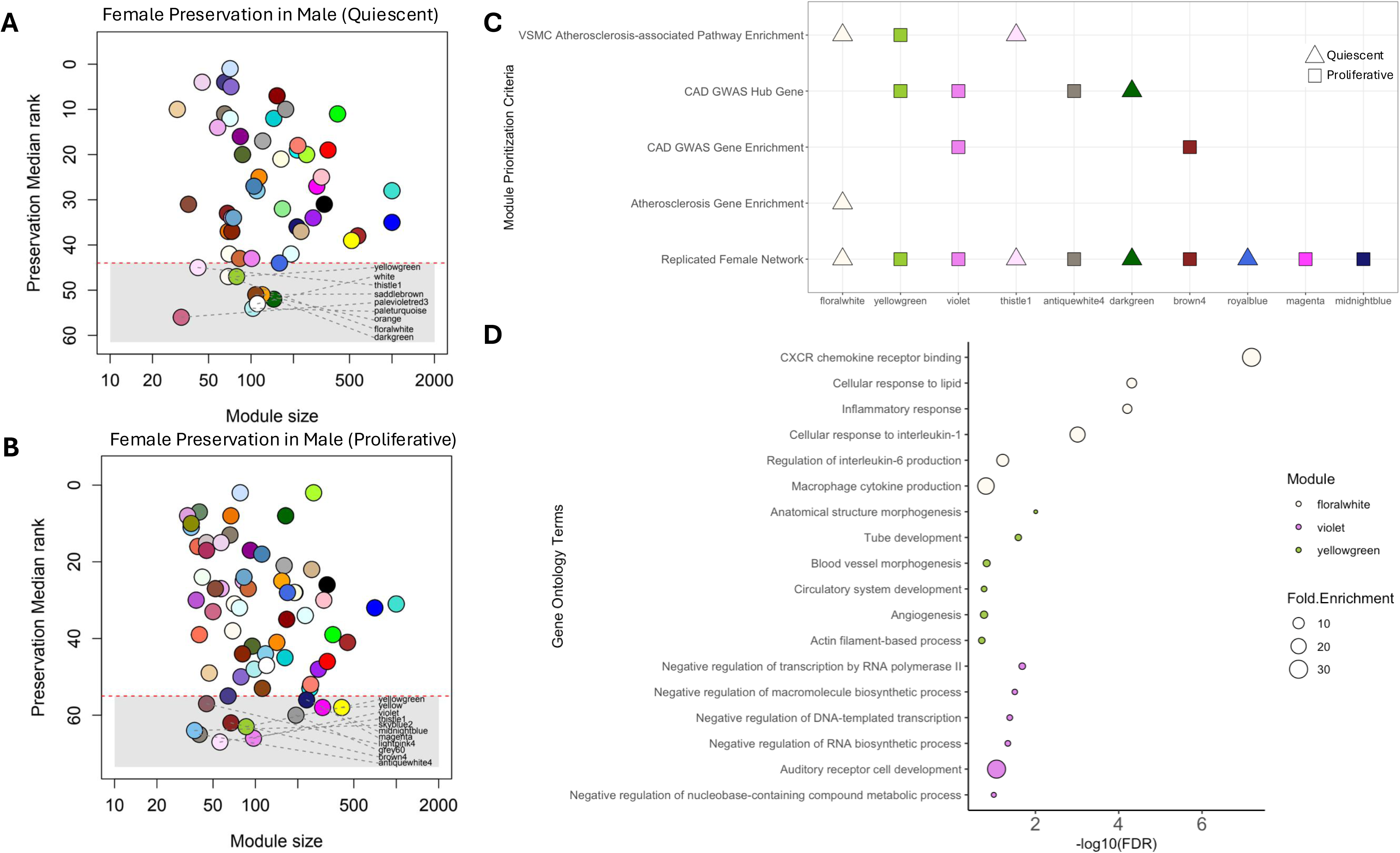
Identification and prioritization of female-specific VSMC coexpression modules associated with atherosclerosis. (A) medianRank scores to evaluate the preservation of female quiescent VSMC modules identified in male quiescent VSMC modules. (B) medianRank scores to evaluate the preservation of female proliferative VSMC modules identified in male proliferative VSMC modules. The red line denotes the bottom 20^th^ percentile of preservation scores. Modules at or below the red line represent the least preserved modules. (C) Prioritization of unpreserved female modules based on replication in previously identified female-biased modules, enrichment for atherosclerosis-associated genes, enrichment for CAD GWAS genes, the presence of CAD GWAS hub genes, and the enrichment of VSMC atherosclerosis-associated Gene Ontology Terminology (GO Term) pathways (Biological Process (BP) and Molecular Function (MF)). (D) GO Term enrichment (BP and MF) of the top three prioritized modules in 2C (floralwhite, yellowgreen, and violet). Each point is scaled according to fold enrichment values.

We aimed to replicate the unpreserved female modules in an independent dataset. A previous study of bulk RNAseq from carotid atherosclerotic plaques from 158 female and male age-matched donors in the Athero-Express cohort identified seven female-biased modules^10^. We compared the module membership in each of our 22 modules to the module membership of each of their seven modules using Fisher’s Exact test. We identified that 10 of the 22 (xx%) unpreserved modules in our dataset were enriched for female-biased networks identified in the female human carotid plaques from the Athero-Express cohort (**Supplemental Table 4**). To prioritize the ten replicated modules for disease relevance, we performed additional filtering steps.

First, we identified which of the 10 GRNs were relevant to genetic susceptibility to CAD. Genome-wide association studies (GWASs) have shed light on the genetic architecture of CAD susceptibility; therefore, we manually curated a list of genes found in loci associated with CAD in GWASs^33–36,56^ (**Supplemental Table 5**). We found that two modules were enriched for CAD GWAS genes (**Figure 2C**). Second, we identified which of the 10 GRNs were relevant to atherosclerosis development. We utilized a curated list of atherosclerosis genes obtained from DisGenet^57^ (C0004153, C0155626, C0151744) and Ingenuity Pathway Analysis^9^ (**Supplemental Table 5**). We found that one module was enriched for known atherosclerosis genes (**Figure 2C)**. Third, we assessed the enrichment of GO Term gene sets (Biological Process and Molecular Function) for each GRN (**Supplemental Tables 6,7**). We found that three modules were enriched for pathways relevant to SMCs and atherosclerosis (FDR ≤ 0.25). GRN_floralwhite_ was enriched for inflammatory and immune response pathways^19^. GRN_yellowgreen_ was enriched for actin filament-based and vascular remodeling processes^58^. Lastly, GRN_thistle1_ was enriched for various MAPK signaling pathways^59^ (**Figure 2C)**. Finally, we identified which replicated GRNs had CAD GWAS genes as hub genes. Hub genes are highly connected genes within a GRN that are central to the module’s network structure and are predicted to play key roles in the biological processes associated with the module^60^. CAD GWAS hub genes could serve as critical nodes in disease-associated regulatory networks. We identified four GRNs meeting this criterion (**Figure 2C, Supplemental Table 8**). The GRNs with the highest prioritization score were GRN_floralwhite_, GRN_yellowgreen_, and GRN_violet_ (Figure 2C). However, only GRN_floralwhite_ and GRN_yellowgreen_ linked disease-associated genes with biological processes directly related to the progression of atherosclerosis in the vascular wall **(Figure 2D**).

The final step in characterizing these two GRNs involved exploring the associations between sex-specific gene expression and atherosclerosis-related phenotypic data. We used the same cohort of 151 SMCs to measure rates of 12 cellular phenotypic associations incorporating VSMC proliferation, migration, and calcification^17^. For GRN_floralwhite_, we first conducted a two-way ANOVA analysis to assess the impact of sex and gene expression across all 12 phenotypes and identified osteogenic calcification to have the highest number of significant associations (p ≤ 0.05, **Supplemental Table 9**). We then identified significant associations between female gene expression and osteogenic calcification and found 12 genes associated – all capturing female-specific correlations (p ≤ 0.05, **Figure 3A**). In GRN_yellowgreen_, we identified proliferation response to TGFβ1 treatment to have the highest number (10) of gene-by-sex associations following two-way ANOVA testing (**Supplemental Table 9**). We identified 24 genes significantly correlated with female gene expression and TGFβ1-induced proliferation. Only two genes were also significantly correlated in males (**Figure 3B**). Both calcification and TGFβ are associated with the development of atherosclerotic plaque^61–63^, suggesting that the two prioritized female-specific GRNs may provide valuable insights into female-specific genetic regulation contributing to pathogenesis.

**Figure 3:**
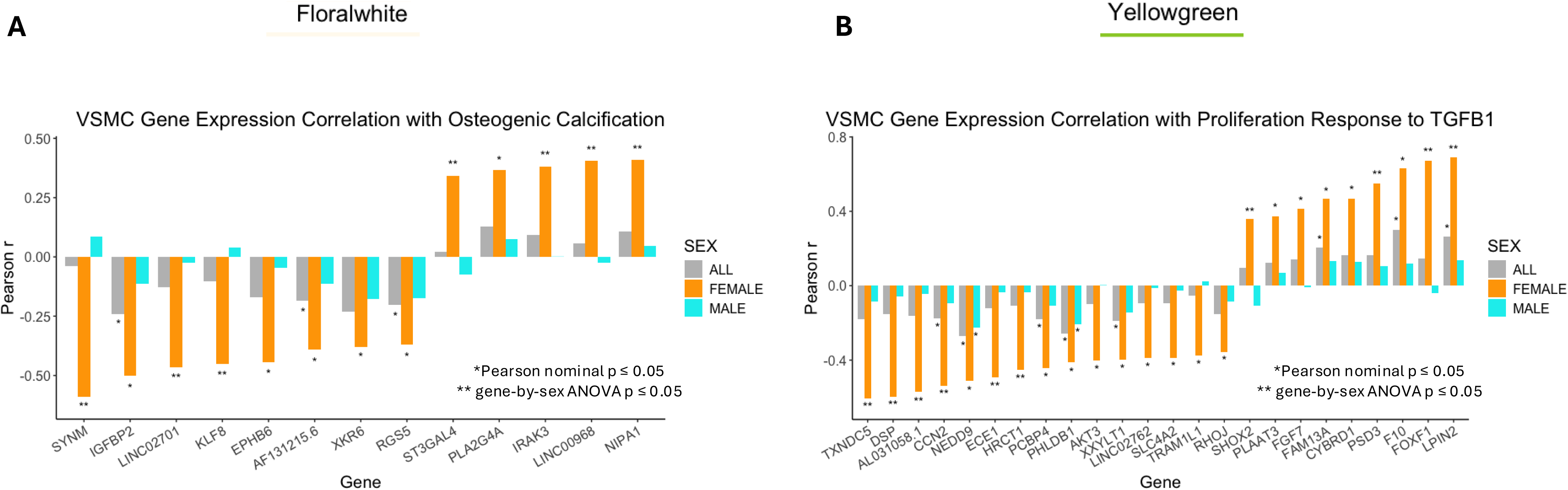
Female-specific gene expression correlations with *in vitro* VSMC atherosclerosis-associated phenotypes. (A) Pearson correlation coefficients (r) for genes in GRN_floralwhite_ with female VSMC gene expression (orange) associations with Osteogenic Calcification cellular phenotype. For each gene, male gene expression (cyan) and combined female and male gene expression (grey) correlations are shown. (B) Pearson correlation coefficients (r) for genes in GRN_yellowgreen_ with female VSMC gene expression (orange) associations with Proliferation Response to TGFβ1. For each gene, male gene expression (cyan) and combined female and male gene expression (grey) correlations are shown. * represents a Pearson nominal p-value ≤ 0.05. ** represents a gene-by-sex two-way ANOVA nominal p-value ≤ 0.05.

### Independent sex-specific evidence for prioritized gene regulatory networks

To study these two prioritized female-biased GRNs in more detail, we used single-cell RNA sequencing data from carotid plaques of 46 patients (20 women and 26 men, **Figure 4A**). This dataset identified 10 cell-type clusters in atherosclerotic plaques, including ACTA2+ SMCs. We first calculated the module score of both GRNs within ACTA2+ SMCs to validate our sex-biased results. The module score is a proxy for the average expression of the genes within each GRN for each cell^64^. We performed this analysis in a sex-stratified manner to identify if there were differences between cell-type clusters and gene expression signatures within each GRN due to sex. For ACTA2+ SMCs, the genes in GRN_yellowgreen_ had a higher module score in females compared to males, whereas the genes in the GRN_floralwhite_ had a lower module score in females compared to males **(Figures 4B,C)**. These results support the fibrous plaque phenotypic contributions hypothesized by the GRN_yellowgreen_ and GRN_floralwhite_ pathway enrichments. We would expect the fibrous-associated GRN_yellowgreen_ to have a higher module score in females and the inflammatory-associated GRN_floralwhite_ to have a higher module score in males.

**Figure 4:**
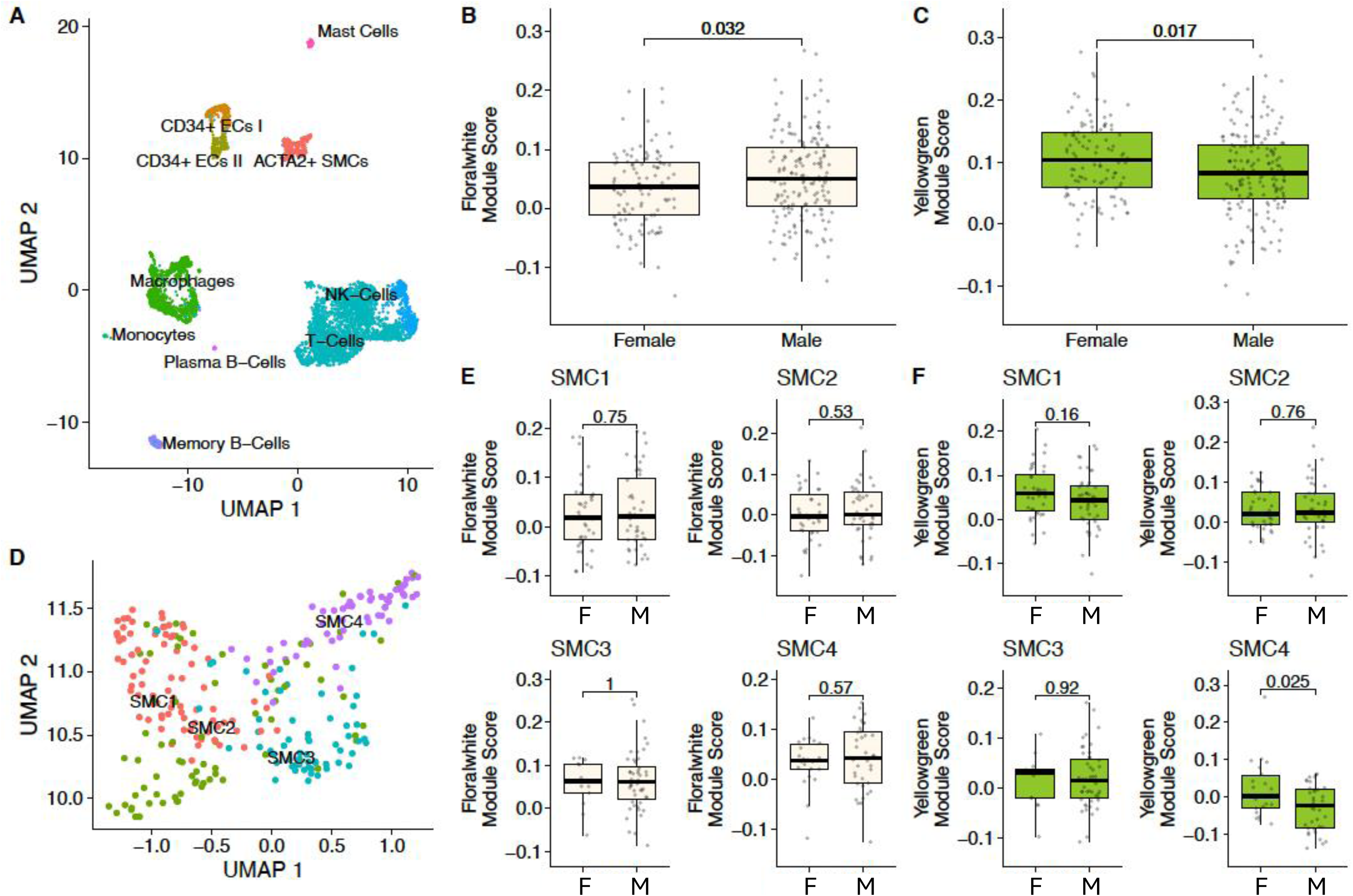
Sex-specific validation of GRN_floralwhite_ and GRN_yellowgreen_ in single-cell RNA-sequencing data of atherosclerotic plaques. (A) UMAP plot of 4,948 single cells from carotid plaque (20 female and 26 male). (B) Module score expression of GRN_floralwhite_ genes in ACTA2+ SMCs comparing female and male single-cell expression data. (C) Module score expression of GRN_yellowgreen_ genes in ACTA2+ SMCs comparing female and male single-cell expression data. (D) Zoom-in UMAP of 672 smooth muscle cells and the defined four subclusters from carotid plaque. (E) Module score expression of GRN_floralwhite_ genes in each of the four SMC subtypes comparing female and male single-cell expression data. (F) Module score expression of GRN_yellowgreen_ genes in each of the four SMC subtypes comparing female and male single-cell expression data.

SMCs are highly plastic cells that undergo phenotypic switching during the progression of atherosclerosis. Identifying SMC phenotypic states using single-cell RNA sequencing is a common method to understand genes representative of SMC dedifferentiation and unique phenotypic contributions^65,66^. Therefore, we reexamined the female and male module scores from each GRN within the four distinct SMC cell states (**Figure 4D**). We did not find differences in module scores between females and males across any SMC cell states for genes within GRN_floralwhite_ (**Figure 4E**). However, we observed a difference between female and male module scores in the SMC4 cluster for genes within the GRN_yellowgreen_ (**Figure 4F**). The module score was higher for females compared to males. SMC4 is enriched for both contractile biological processes and cytoskeleton and vasculature development pathways (**Supplemental Figure 2**), suggestive of a less synthetic and more plaque stabilizing phenotypic state. These analysis results support the female-biased nature of GRN_yellowgreen_ and complementary evidence of the potential phenotypic contributions associated with GRN_yellowgreen_ and a protective fibrous cap.

### Topology analysis of prioritized female-specific yellowgreen gene regulatory network

Coexpression networks are unable to infer the directionality of nodes within a network; therefore, we applied Bayesian network (BN) algorithms^45,67^ to infer directionality and refine regulatory interactions for GRN_yellowgreen_. This network contains the highest degree of evidence linking female-specific gene expression signatures with female-biased plaque phenotypes. We used a hill-climbing score-based algorithm with 10,000 iterations to infer edge weights and directionality for each of the 86 genes within GRN_yellowgreen_. We then trimmed low-weight edges to yield a scale-free topological distribution representative of biological networks where most nodes have few connections while a small number of nodes have many connections^47,68^ (**Figure 5A**, **Supplemental Table 10**). Next, we performed key driver (KD) analysis on BN_yellowgreen_ to identify the genes with the highest regulatory potential that may govern fibrous plaque-associated pathways^48^. We first computed the size of the h-layer neighborhood for each node in the BN_yellowgreen_. We then defined the average size of each h-layer neighborhood to score nodes with the highest regulatory potential according to the network topology. We defined KDs as genes within the top 10% of KD scores, resulting in 6 KD genes for BN_yellowgreen_ (*TMSB15B, MYH9, CAMSAP1, IFFO2, CSRNP1,* and *ARMC5*; **Supplemental Table 11**). These six genes are expected to have a more significant effect in regulating downstream gene expression and the function of biological pathways within the yellowgreen network. To ensure that these KD genes are specific to female networks, we used the expression values for the same genes from the male donors and recreated the BN_yellowgreen_ (**Supplemental Table 10**). We then compared the topology of the male and female networks and identified KDs in each network. The female-specific network had significantly higher mean edge connectivity compared to the male network when using untrimmed edge weights (**Figure 5B**). There were no overlapping KD genes between female and male BN_yellowgreen_ (**Supplemental Table 11**). To further show the sex-specificity of KDs in the female BN_yellowgreen_, we separately compared the number of genes directly downstream of the female KDs in the male and female networks. The six female BN_yellowgreen_ KD genes had 57 downstream edges, one node away in the female network versus only 32 downstream edges, one node away in the male network (**Figure 5C**). The KD gene that showed the largest difference in predicted downstream genes between female and male networks was *MYH9*. We found that the KD designation of *CSRNP1* and *MYH9* was reproduced in female-specific networks in the vascular tissues from CAD (STARNET^9,39^) and stroke (Athero-Express) patient cohorts, respectively (**Figure 5D**). However, only *MYH9* was differentially expressed between males and females in the atherosclerotic aorta of 160 age-matched donors in the STARNET cohort (P_adj_ = 0.03). Therefore, we hypothesized that *MYH9* plays a sex-specific key regulatory role in fibrous-associated plaque progression.

**Figure 5:**
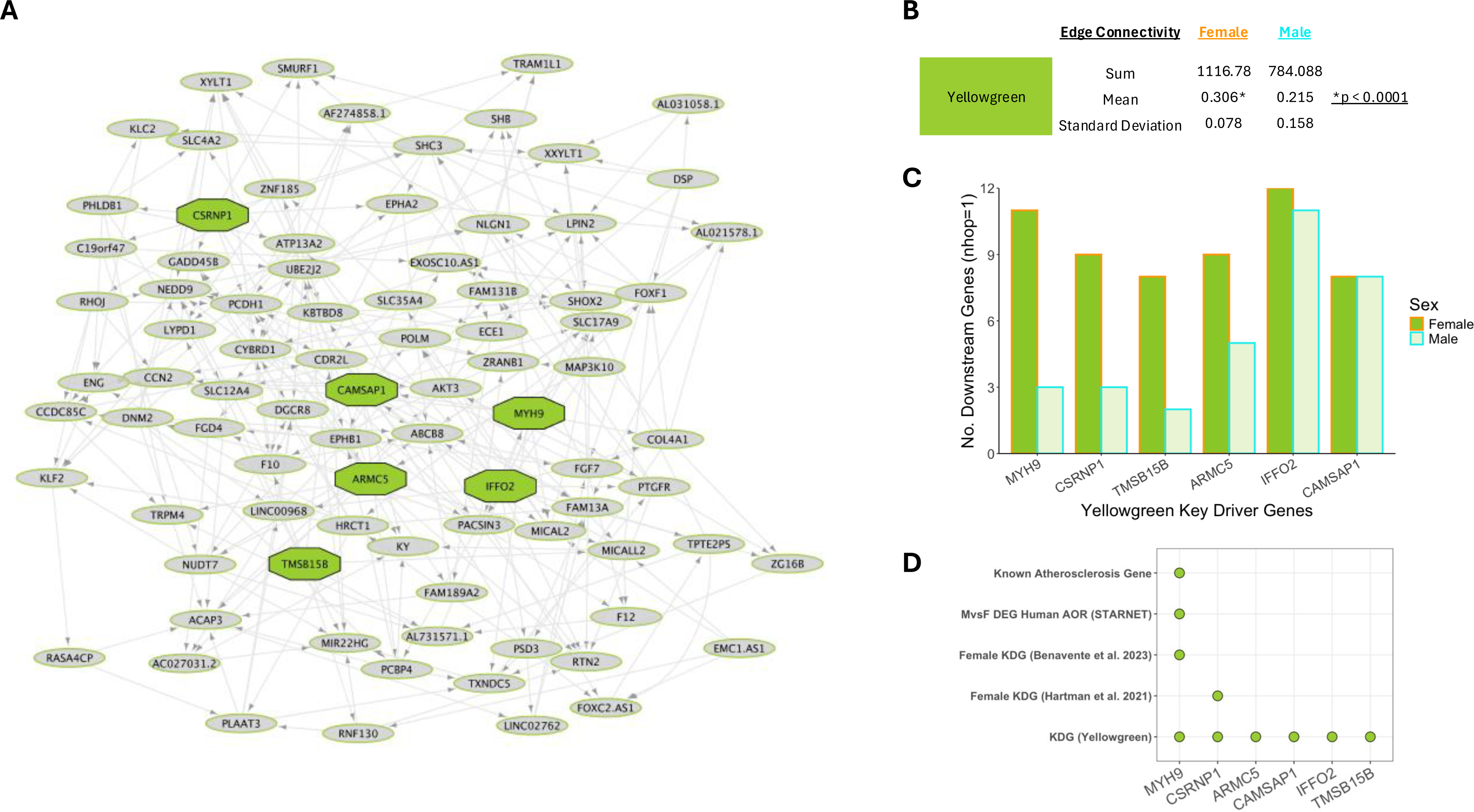
Bayesian network and key driver analysis identifies MYH9 as a female-specific key driver gene of GRN_yellowgreen_. (A) Bayesian network (BN) created from genes in GRN_yellowgreen_ using female gene expression data. Octagonal nodes represent key driver genes. (B) Network topology summary statistics for edge weight characteristics of BN_yellowgreen_ were generated using female gene expression data and BN_yellowgreen_ were generated using male gene expression data. The mean edge connectivity is significantly higher in the female BN versus the male BN (p < 0.001). (C) The number of downstream genes one node away from each of the six key driver genes identified in Figure 5A in both the female and male-generated BNs. (D) Prioritization of female-specific key drivers based on replication in previously identified female-biased modules from the STARNET and Athero-Express cohort, differential gene expression between female and male human aortas with CAD (STARNET), and whether it is a known gene involved in atherosclerosis.

### MYH9 protein level in atherosclerotic plaques is correlated with fibrous-associated biological processes

Transcription is just one layer of regulation and does not always reflect protein expression^69,70^. Therefore, to better understand how biological processes and potential disease mechanisms are regulated by MYH9, we incorporated untargeted LC-MS proteomics data from 200 patients (149 male, 51 female) undergoing carotid artery endarterectomy^38^ (see Methods). Of the 1499 proteins measured, 10 overlapped with genes in GRN_yellowgreen_. We identified proteins differentially abundant across sexes (**Supplemental Table 12**) and found MYH9 protein to have the most significant difference, upregulated in female plaques (logFC = 0.4, p_adj_ = 0.015, **Supplemental Figure 3A**). Due to the low protein gene overlap within GRN_yellowgreen_, to predict MYH9 functional interactions within the proteomic data, we expanded the analysis to all 1499 proteins identified within the plaque samples and first calculated the correlations of all proteins with MYH9 using female and male samples independently. GO Term analysis revealed an enrichment of actin filament and cytoskeleton processes only in proteins significantly correlated with MYH9 in female samples (**Supplemental Figure 3B,C**). When combining female and male data, we identified 642 positively and 133 negatively correlated proteins (p_adj_ ≤ 0.05; **Supplemental Table 13**). GO Term analysis of positively correlated proteins showed enrichments for actin-filament organization, cell-matrix adhesion, and extracellular matrix organization, capturing similar biological processes as GRN_yellowgreen_ and SMC4 subcluster (**Supplemental Figure 4A**). Proteins negatively correlated were enriched primarily for a variety of immune response pathways (**Supplemental Figure 4B**). Upregulation of MYH9 in female plaques points to a potential regulatory mechanism associated with modulated ECM content and inflammatory signaling recapitulating the stable, fibrous plaque phenotype captured at the transcriptomic level.

### MYH9 protein level in atherosclerotic plaques is associated with fibrous plaque characteristics

To test the hypothesis that MYH9 is associated with fibrous, plaque erosion phenotypes in females, we investigated the relationship between MYH9 and carotid plaque characteristics assessed in 200 patients from the Athero-Express biobank. The carotid plaques in this biobank have been characterized for SMC content using ɑ-actin staining and lipid content using H&E and picrosirius red stains^21,71^. We first calculated the association of MYH9 protein with male and female plaque characteristics. There was a positive correlation between MYH9 protein levels and SMC content in males and a near-significant correlation in females, likely due to differences in sample size (149 male vs 51 female samples) (**Figure 6A**). When the samples were combined, SMC content associations within the plaque and levels of MYH9 were significant (p = 0.0076, **Supplemental Figure 5A**). For associations with lipid content in plaque, we found a sex-specific association. Increased levels of MYH9 were associated with lower levels of lipid content only in females (p = 0.015, **Figure 6B, Supplemental Figure 5B**). Together, these results suggest that increased levels of MYH9 are associated with higher amounts of SMCs and lower amounts of lipid, suggesting an association with fibrous plaques more representative of a plaque erosion phenotype. The carotid plaques in this biobank were further classified based on plaque vulnerability and overall cellular composition to define asymptomatic versus symptomatic plaques as well as fibrous versus atheromatous plaques. Therefore, we investigated the difference in MYH9 abundance in symptomatic and asymptomatic plaques and observed that MYH9 expression was higher in asymptomatic plaques (p = 0.02, **Supplemental Figure 5C**). We also investigated the relationship between MYH9 abundance and plaque classification. We found a female-specific association between elevated MYH9 levels and fibrous plaques versus atheromatous plaques (p = 0.01, **Figure 6D**). This suggests that MYH9 may play a distinct and critical role in the development of fibrous plaques in females, potentially contributing to plaque stability and the erosion phenotype commonly observed in females.

**Figure 6:**
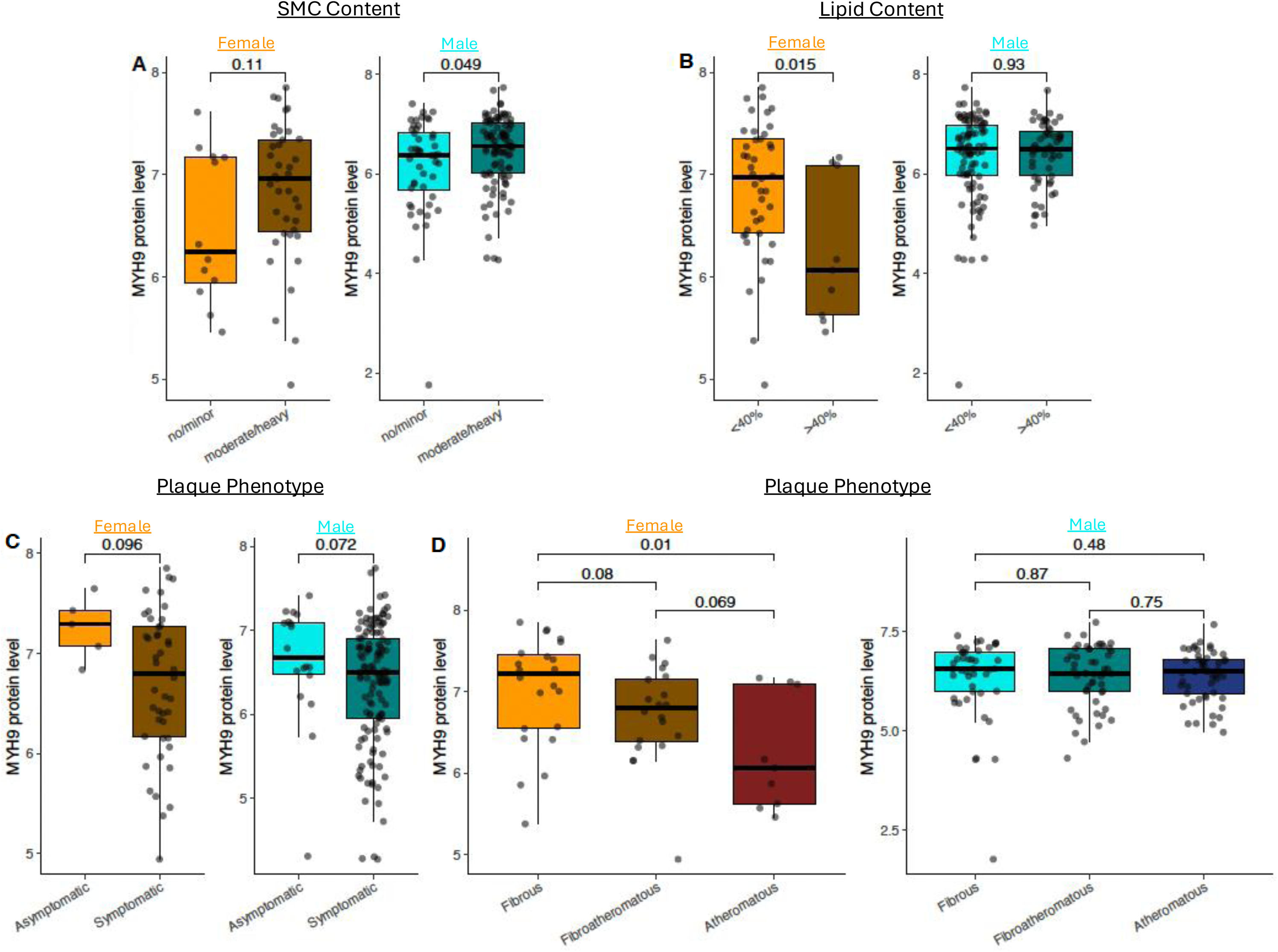
MYH9 protein associations with carotid plaque characteristics suggest both a protective and female-specific role. (A) MYH9 protein levels for plaques defined to have no/minor SMC content and moderate/heavy SMC content in females (orange, p = 0.11) and males (cyan, p = 0.049). (B) MYH9 protein levels for plaques with a lipid core less than 40% and greater than 40% of the total plaque area in females (p = 0.015) and males (p = 0.93). (C) MYH9 protein levels for plaques defined to be asymptomatic and symptomatic in females (p = 0.096) and males (p = 0.072). (D) MYH9 protein levels for plaques defined to be fibrous, fibroatheromatous, and atheromatous in females (p = 0.01, p = 0.069, p = 0.08) and males (p = 0.48, p = 0.75, p = 0.87).

## Discussion

Sex differences in atherosclerosis are often complex and not fully understood. While it is clear that males and females exhibit differences in plaque morphology, with females typically having more stable fibrous plaques and males more vulnerable, atheromatous plaques, the underlying molecular mechanisms remain unclear^2^. The lack of female-centric studies often leads to assumptions based on male-dominated data that may not fully reflect the pathophysiology of females^72^. This bias is gradually being addressed, but current methods still possess significant limitations. Systems biology approaches are common methods for studying complex diseases such as atherosclerosis^18^. Recent studies investigated how GRNs generated from plaque tissue RNA sequencing data differ between males and females in the development and presentation of atherosclerosis. These efforts highlight how gene expression and network connectivity are heavily influenced by sex. Sex differences in tissue composition of atherosclerotic plaques have been described before^73^. Therefore, comparisons of RNA sequencing data obtained from plaques may be confounded by cellular composition differences rather than gene expression levels in each cell type found in plaques. SMC content is a key deterministic feature distinguishing thin versus thick fibrous caps that influence plaque erosion and plaque rupture phenotypes^20^. Sex-stratified SMC GRNs can better detect the sex differences that exist on a cellular level to unravel molecular driving forces describing the higher prevalence of fibrous plaques in females.

In this study, we utilized RNA sequencing data from human VSMCs isolated from male and female heart transplant donors, allowing us to construct sex-specific GRNs. These GRNs were further analyzed for preservation across sexes to identify female-biased network structures. Preservation statistics between GRNs can identify unique aspects of differential conditions that methods such as differential expression analysis and gene set enrichment analysis are unable to capture. Within this cohort of VSMCs, only 8 genes were shown to be differentially expressed between males and females^74^. On the other hand, our method of characterizing the differences in the topology of GRNs identified female-biased gene connectivity enriched for inflammatory pathways and vascular remodeling processes and associated with key atherosclerosis-relevant phenotypes of calcification and TGFβ -induced proliferation. Each of these phenotypes are associated with sex-specific mechanisms of plaque development. Females have fewer coronary calcifications leading to more stable lesions^75^, and TGFβ signaling responses are linked to sex hormone-specific regulation^16^. We were able to reproduce the female-biased GRN results within single-cell RNAseq data. This analysis added a critical layer of validation to our approach and represents the power of utilizing network preservation statistics to identify sex differences. The single-cell RNAseq data also supported the fibrous plaque-association hypothesis of GRN_yellowgreen_. Further analysis of GRN_yellowgreen_ using BN methods and proteomics allowed us to detect MYH9 as the key driver predicted to regulate actin, ECM, and vascular remodeling processes, potentially influencing the progression of fibrous plaques in females. We tested this hypothesis within carotid plaques and found MYH9 to be associated with an increased prevalence of fibrous plaques over atheromatous plaques only in females. MYH9 is involved in actin filament dynamics, which is critical for cellular processes such as motility, morphogenesis, and contractility^76,77^. Increased MYH9 expression in female VSMCs may enhance the contractile phenotype and limit excessive migration and proliferation during disease progression, promoting a more stable plaque formation. Our results provide important insights into sex-specific gene expression patterns that could inform our understanding of the molecular underpinnings of fibrous plaque formation, plaque erosion, and the broader pathophysiology of atherosclerosis in males and females.

Deciphering the exact mechanisms of sex-specific contributions to atherosclerosis phenotypes associated with MYH9 requires further investigation. Sex hormones and chromosomes likely interact in complex ways to influence cardiovascular disease risk^78^. Sex chromosomes contribute to sex-specific gene expression through mechanisms such as X-inactivation, where certain genes escape inactivation on the X chromosome, leading to differential expression in females compared to males^79^. Protein correlations for MYH9 revealed RPS4X as the highest correlated protein in both sexes. *RPS4X*, an X-linked gene known to regulate translation^80^, has been shown to escape X-inactivation in females and may help explain the differential protein regulation of MYH9 between females and males^79^. On the other hand, sex hormones could also play key mechanistic roles in the development of atherosclerotic plaque between sexes.

Estrogen, for instance, has been linked to a protective effect on the vascular wall by inhibiting the proliferation and migration of VSMCs through MAPK/ERK and PI3K/AKT pathways delaying the formation and development of plaques^81^. It has also been shown to enhance cholesterol efflux and reduce foam cell formation derived from VSMCs^82^. This could potentially describe the contractile and more fibrous phenotype associated with GRN_yellowgreen_ as well as the female-specific association between MYH9 and fat content within plaques. The interplay between sex chromosomes and sex hormones likely creates a multifaceted regulatory environment that shapes atherosclerotic disease progression in a sex-specific manner. Therefore, understanding both the chromosomal and hormonal influences on gene regulation in vascular cells, particularly in the context of MYH9 and other key atherosclerosis-related genes, will be essential to uncover the full scope of sex differences in cardiovascular disease.

Although this study provides compelling evidence for sex-specific differences in the molecular mechanisms underlying atherosclerotic plaque phenotypes, quantifying VSMC phenotypes in cell culture relevant to atherosclerosis has inevitable limitations. For example, the culture conditions lack the key interactions with other cell types and environmental conditions in the vessel wall and atherosclerotic plaque. In addition, atherosclerosis takes decades to develop; therefore, it cannot be adequately replicated *in vitro*. Despite these challenges, cultured human artery SMCs have been successfully used in previous studies to investigate genetic determinants of CAD^83–85^. Due to the difficult nature of capturing vascular wall phenotypes in cellular detail in the arteries of humans, our approach provides a reasonable proxy for in vivo characteristics of VSMCs. In addition, the sample size for female donors was relatively small (n=32), which may limit the generalizability of our findings. However, this limitation guided many of our efforts to repeatedly show that the female-specific GRNs were, in fact, female-specific and atherosclerotic relevant, potentially providing even more confidence in our major findings. This highlights the need for incorporating appropriate sample sizes for females in large cohort studies. To our knowledge, this work is the first to propose a comprehensive approach exploring gene regulatory networks observed in female VSMCs that are not preserved in male VSMCs.

There may be male-specific contributions to plaque rupture phenotypes that we did not investigate. Even though the primary focus of this study was to better understand plaque erosion phenotypes through the investigation of female VSMCs, the results provide a reason to utilize male VSMCs to elucidate mechanisms of plaque rupture phenotypes. Our findings should be considered under the current limitations of employing systems genetics and network analyses. There are many decisions made during the model-building phase and prioritization of GRNs that can affect the results and conclusions.

Our study sheds light on the critical role of sex-specific GRNs in vascular smooth muscle cell biology, specifically focusing on fibrous plaque formation and plaque erosion. By leveraging a multi-dimensional approach, we hypothesize that MYH9 is a key driver of actin and vascular remodeling processes that regulate SMC-rich fibrous plaques. Further investigation is needed into the exact mechanisms of disease and the potential relationship between MYH9 and TGFβ in creating more stable, fibrous caps prone to plaque erosion. Additional research is also necessary to address the limitations of *in vitro* models, small sample sizes, and potential sex-specific contributions to other plaque phenotypes, such as plaque rupture. More in-depth investigations into the interactions between sex chromosomes, sex hormones, and gene regulatory networks will enhance our understanding of how these factors collectively shape cardiovascular disease progression. Ultimately, a better understanding of sex-specific molecular mechanisms in atherosclerosis will pave the way for more targeted therapies and personalized approaches in the prevention and treatment of cardiovascular disease.

### Sources of Funding

Research reported in this publication was supported by the National Institute of Health under Award Numbers F31HL165772 (R.N.P.), R01HI156120 (M.C.), R01HL164577 (J.L.M.B), R01HL148167 (J.L.M.B.), R01HL148239 (J.L.M.B.), R01HL166428 (J.L.M.B.), and R01HL168174 (J.L.M.B.), an American Heart Association Established Investigator Award 24EIA125067 (M.C.), and the Leducq AtheroGEN consortium 22CVD04 (to M.C., H.d.R., and J.B.). J.L.M.B also acknowledges support from the Swedish Research Council (2018-02529 and 2022-00734), the Swedish Heart Lung Foundation (2017-0265 and 2020-0207), and the Leducq PlaqOmics(18CVD02) consortia. MdW acknowledges support from from ZonMW (ZonMW Open competition 09120011910025), NWO (Epi-Guide-Edit - KICH1.ST01.20.045) and the European Union (Horizon Europe research and innovation program under Marie Sklodowska-Curie Actions Doctoral Network program 2022 under Grant Agreement No. 101119370; MIRACLE).

