## Supplemental Figures for "Female-biased vascular smooth muscle cell gene regulatory networks predict MYH9 as a key regulator of fibrous plaque phenotype"

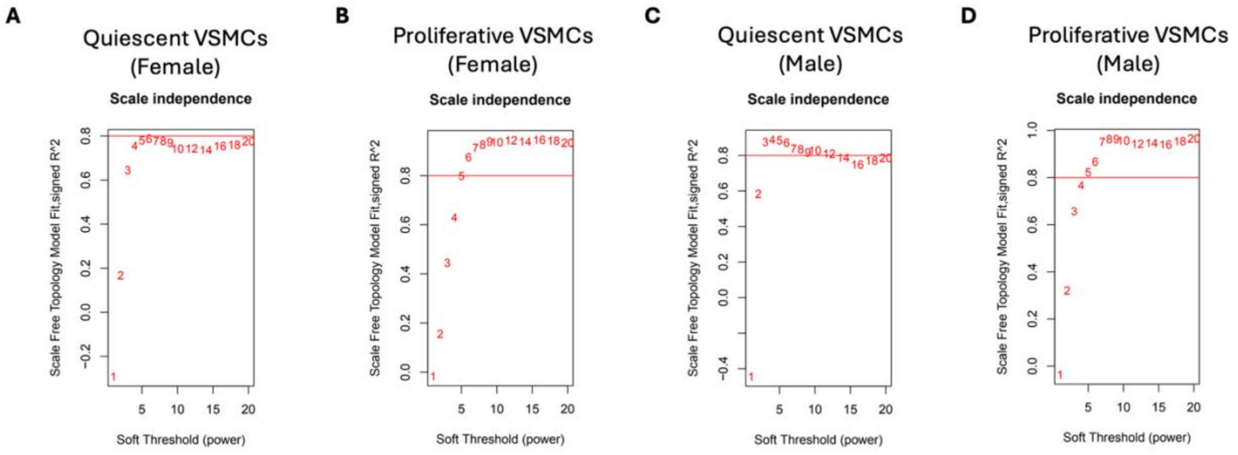


**Supplemental Figure 1:** Determination of soft-thresholding power in Weighted Gene Co-expression Network Analysis. The scale-free fit index (y-axis) as a function of the soft-thresholding power (x-axis) for gene expression of (A) female quiescent, (B) female proliferative, (C) male quiescent, and (D) male proliferative vascular smooth muscle cells.

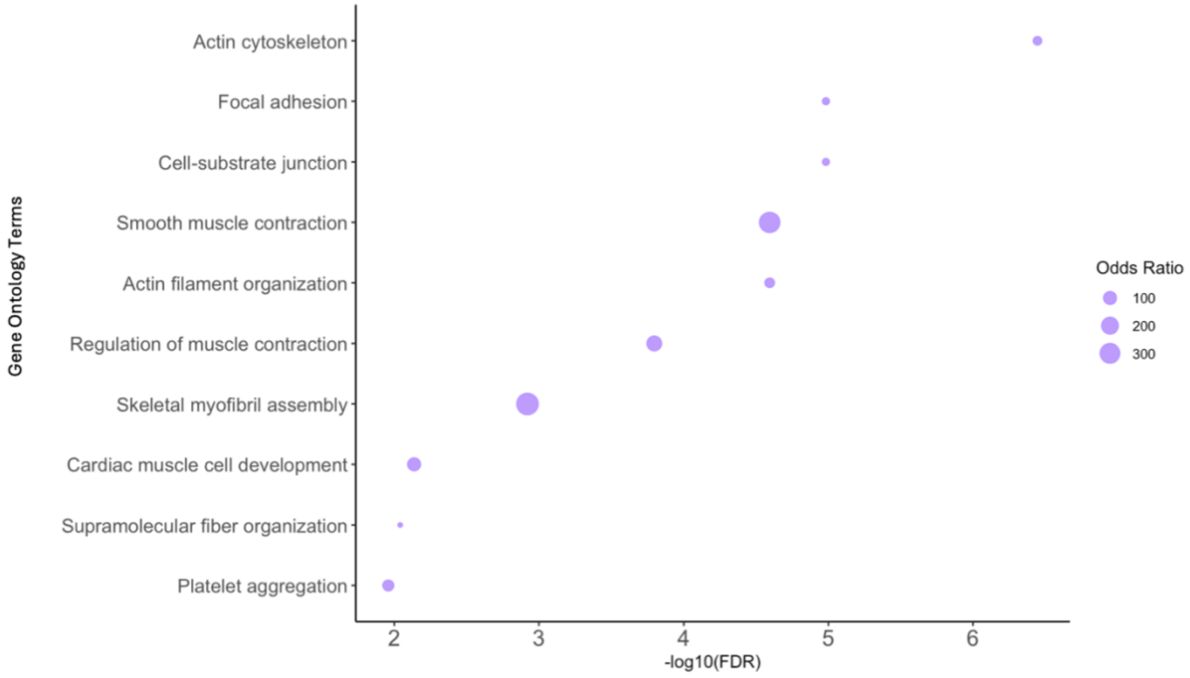


**Supplemental Figure 2:** Top 10 enriched Gene Ontology terms from differentially expressed genes of SMC4 subtype in carotid scRNAseq.

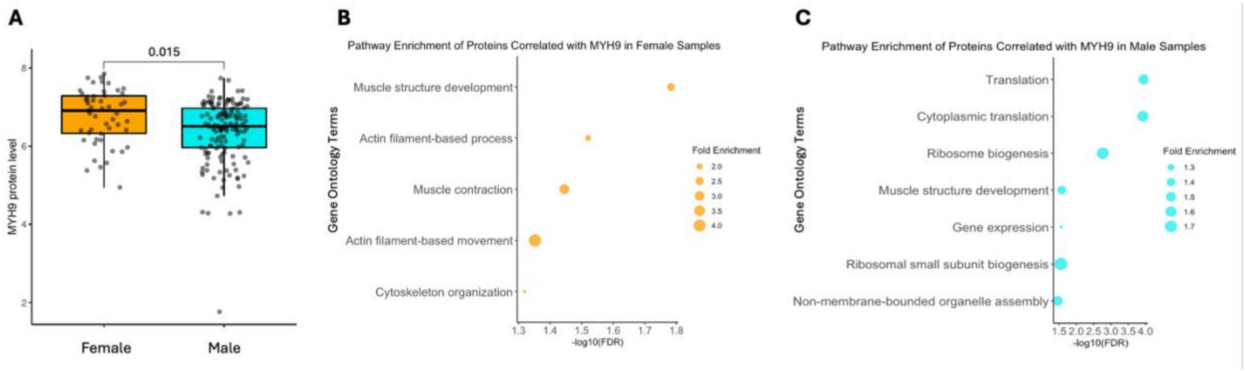


**Supplemental Figure 3:** (A) Differential abundance of MYH9 protein levels in carotid plaques of 51 females and 149 males from the Athero-Express cohort (p = 0.015). Biological Process Gene Ontology enrichment of (B) proteins correlated ($p_{adj} \leq 0.05$) with MYH9 in female samples only (n=51) and (C) proteins correlated ($p_{adj} \leq 0.05$) with MYH9 in male samples only (n=149).

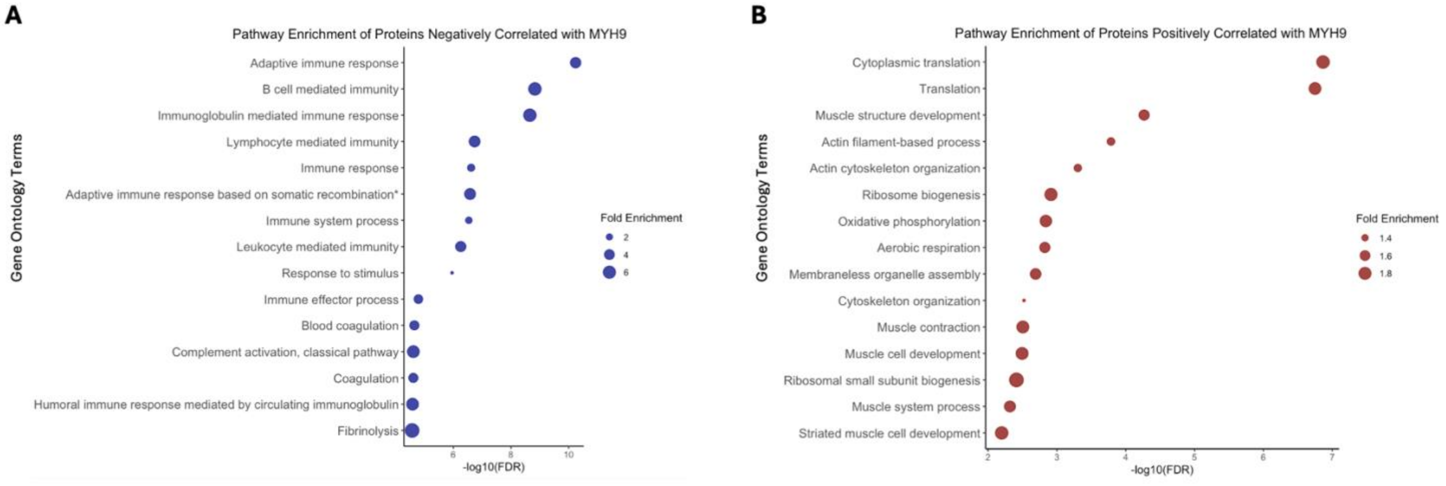


**Supplemental Figure 4:** Biological Process Gene Ontology enrichment of (A) proteins negatively correlated with MYH9 and (B) proteins positively correlated with MYH9 ($p_{adj} \leq 0.05$) in carotid plaques of 200 female and male combined samples.

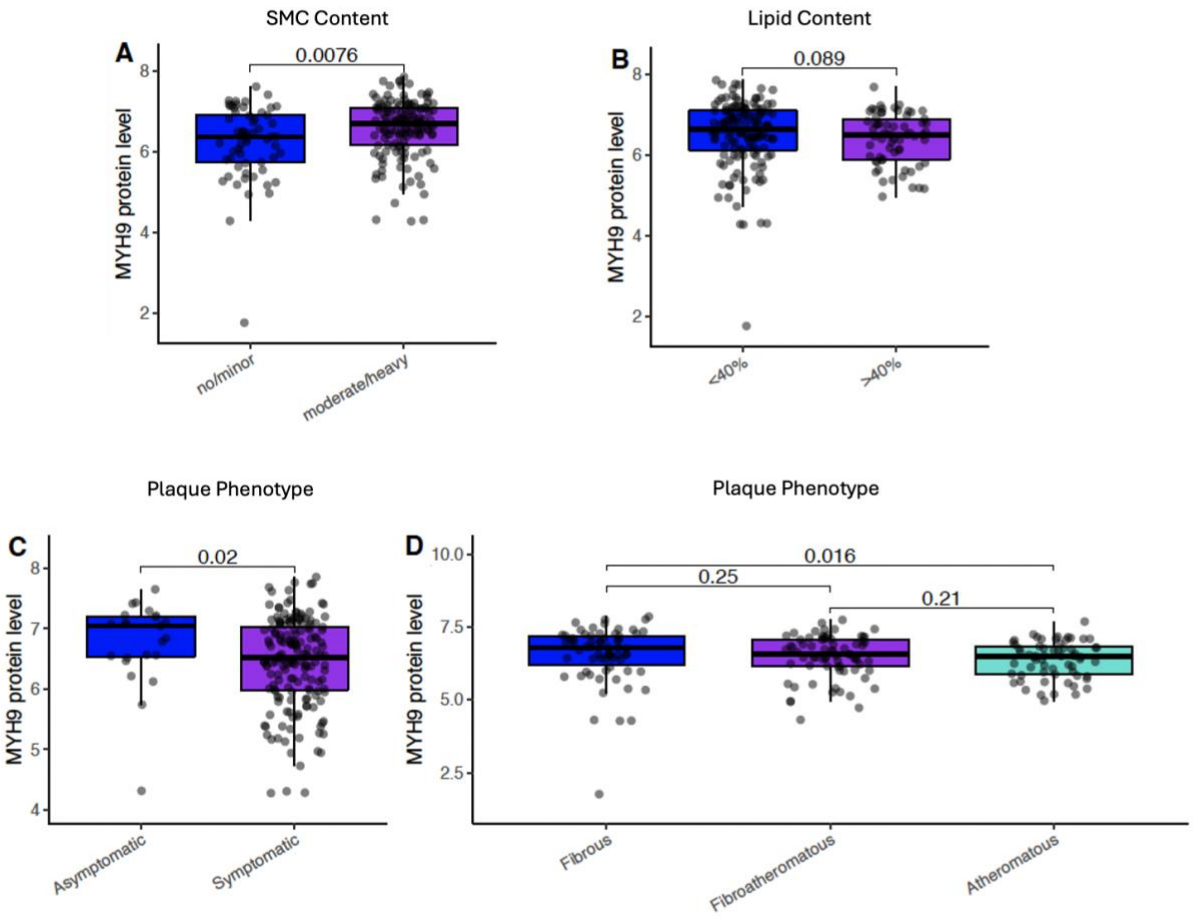


**Supplemental Figure 5:** (A) MYH9 protein levels for plaques defined to have no/minor SMC content and moderate/heavy SMC content in both female and male samples combined ($p = 0.0076$). (B) MYH9 protein levels for plaques with a lipid core less than 40% and greater than 40% of the total plaque area in both female and male samples combined ($p = 0.089$). (C) MYH9 protein levels for plaques defined to be asymptomatic and symptomatic in both female and male samples combined ($p = 0.02$). (D) MYH9 protein levels for plaques defined to be fibrous, fibroatheromatous, and atheromatous in both female and male samples combined ($p = 0.016$, $p = 0.21$, $p = 0.25$).
